# Resolving the Deep Sleep Dual Indeterminacy Problem: Context-Dependent Slow-Wave Activity Modeling Predicts Neurobehavioral Fatigue Where Clinical Sleep Modeling Fails

**DOI:** 10.64898/2026.03.25.714331

**Authors:** Shashaank Vattikuti, Hua Xie, Carson C Chow, Thomas J Balkin, John D Hughes

**Affiliations:** TrekMH, Reston, VA; Laboratory of Biological Modeling, NIDDK, NIH, Bethesda, MD; Walter Reed Army Institute of Research, Silver Spring, MD

## Abstract

Deep sleep is widely considered to be the most recuperative component of sleep restoration. Accordingly, a positive relationship between naturally occurring deep sleep and function (e.g., cognitive performance) is often assumed. However, this assumption warrants closer examination—particularly given the rise of sleep tracking that emphasizes traditional sleep metrics and their implied predictive value. We present evidence that while clinical deep sleep scoring provides no predictive value, slow-wave activity (SWA) exhibits a paradoxical association with both improved and worsened neurobehavioral fatigue following sleep deprivation. Specifically, we found that SWA-based models account for approximately 50–60% of the inter-individual variance in recovery from sleep deprivation. Remarkably, when regressed against recovery from sleep deprivation, SWA during the baseline sleep night showed a negative association (normalized β = (–)0.5, p = 0.001) while in the same model SWA during the subsequent wakefulness period showed an opposite positive association (normalized β = 0.5, p = 0.001). Furthermore, although the group-averaged SWA while behaviorally awake increased with impairment across the sleep deprivation period, individual-level data revealed an inverse relationship: individuals more resilient to sleep deprivation exhibited greater SWA in-between mental test sessions and less corresponding impairment during wakefulness suggestive of a protective effect. These findings identify a Deep Sleep Dual Indeterminacy Problem — simultaneous measurement and causal indeterminacy — that explains why clinical sleep staging fails as a functional biomarker across a wide range of outcomes, and provide a principled framework for next-generation sleep metrics grounded in continuous electrophysiology and temporal modeling.

## Introduction

Deep sleep is a critical component of cognitive restoration^1−3^. As a result, a positive relationship between measured duration of deep sleep — specifically as codified by clinical scoring (AASM N3) — and functional outcomes is often assumed. This assumption merits renewed scrutiny, particularly in light of the recent proliferation of consumer sleep trackers that target clinical sleep metrics. Their widespread adoption has conferred unwarranted social credibility on these measures, further amplified by large-scale population studies that, by virtue of enormous sample sizes, lend statistical weight to weak effects while obscuring fundamental interpretation issues^4^. While an obvious concern among specialists has been the accuracy of devices and algorithms against the clinical gold-standard polysomnography (PSG)^5^, these efforts have largely ignored a more fundamental issue: the construct validity of the AASM scoring model itself. Consumer sleep trackers have nonetheless popularized clinical sleep scores as accurate and interpretable biomarkers of brain restoration or, more generally, health^4−7^. This uncritical acceptance has already moved these metrics into actionable tools — directly, and more consequentially, as unsupported composite ‘functional’ scores^6^ and AI health coaching pipelines built on weak evidence and expert opinion^7^.

Indeed, when subjective sleep quality is predicted from the gold-standard PSG, findings are inconsistent across studies and notably, N3 does not emerge as a reliable contributor in any of them^8−11^. Some evidence even suggests N2 — labeled by consumer trackers as ‘light sleep’ and implicitly poor sleep — may in fact be a positive correlate of subjective sleep quality, while N1, lumped together with it, shows the opposite relationship^9^. In one study, N3 was even associated with the lightest subjective sleep experience^12^. While these studies used subjective outcomes, this lack of robust and consistent associations between sleep scoring and functional outcomes extends to objective performance measures as well. For example, the neurobehavioral gold-standard for fatigue is the Psychomotor Vigilance Test (PVT) — a test uniquely sensitive to acute and chronic homeostatic effects and circadian modulation^13’14^. No studies have demonstrated a clear positive association between clinical sleep stages including N3 and subsequent PVT performance. In a prior study from our group, Vattikuti et al. demonstrated this lack of correlation with conventional sleep scoring but found that when focusing on sleep microarchitecture, a strong functional signal emerged — specifically in REM neural activity^15^. The findings supported the hypothesis that clinical sleep scoring is a highly lossy compression of the underlying electrophysiology — the functionally relevant information resides in the raw signal but is discarded by the staging process. Here we test whether a parallel signal exists in deep sleep, and whether the resulting framework can explain why clinical staging fails as a functional biomarker — with implications for the development of next-generation sleep metrics grounded in continuous electrophysiology and temporal modeling.

We argue that operationalizing deep sleep as a predictor is more complex than a simple measurement problem — it is simultaneously a measurement and a causal modeling problem. We term this the *Deep Sleep Dual Measurement and Causal Indeterminacy Problem* — the former reflecting the measurement indeterminacy introduced by discrete sleep staging, which discards the continuous electrophysiological signal, and the latter capturing the reverse causation inherent in a variable that conflates recovery with the homeostatic drive for recovery. This duality stems directly from the foundational work of Borbély and colleagues. Borbély’s pioneering two-process model established that deep sleep increases following sleep deprivation, reflecting a homeostatic rebound — meaning elevated deep sleep may index the drive for recovery as much as recovery itself. Thus, elevated deep sleep may represent better recovery, a response to greater neurophysiological demand (sleep debt), or both. Borbély himself further cautioned against over-reliance on sleep staging, arguing that NREM stages represent a crude and arbitrary subdivision of a continuous process^1^.

Addressing this gap, the present study uses data from a controlled sleep manipulation paradigm. We leverage clinical-grade PSG with expert scoring from a professional sleep laboratory unit along with the PVT to investigate the nuanced relationships between deep sleep physiology and objective fatigue, investigating standard manual polysomnographic scoring but also slow-wave activity (SWA). SWA is a key feature of deep sleep regarded as the primary restorative component3. By comparing the predictive value of standard sleep staging vs. quantitative SWA measures, our goal is to explicate the extent to which these measures relate to functional outcomes – a comparison that has implications for the interpretation of sleep “scores”, and that informs improved quantitative assessments.

We conducted secondary analyses on data originally collected for a non-invasive slow oscillation (SO) (closely related to SWA) stimulation study. This dataset was well-suited for our purposes, as it included ∼45 hours of sleep deprivation recordings across several healthy young adult sleepers, along with baseline and recovery physiology and performance measures. Although neural stimulation was part of the original design, we treated its effects as noise for the current analysis. There are three possible effects: 1) increased induced-SO, 2) reduced SO due to sleep disruption, or 3) no effect. This can be treated as noise, a random variable, potentially reducing the effect size of our observed variables (predictors).

In addition to clinical sleep scores, we examined the full continuum of SWA throughout behavioral sleep (agnostic to sleep scores) and throughout behavioral wake periods. “Behavioral” in this context refers to whether the subject is outwardly engaged or disengaged from its environment. In contrast, neurophysiological sleep processes are now recognized to occur, at least in part, across both behavioral sleep and wake and represent local processes^16’17^. For example, prior human studies have also shown an increase in SWA during prolonged wakefulness, though occasionally at higher spectral frequencies^18−20^.

First, we confirmed that the key clinical and research standard variables changed as expected with the experimentally manipulated sleep conditions. Then, as a benchmark, we investigated the relationship between clinically scored deep sleep (N3) and subsequent vigor (inverse of fatigue) as measured using the PVT. This was evaluated under several sleep-wake conditions. Having failed to find any relationship with sleep parameters based on clinical sleep scoring, we performed more detailed investigations of SWA and vigor and consistently found both strong positive and negative associations.

## Results

### Clinically-Scored Deep Sleep: A Null Result Across All Operationalizations

We first examined whether clinically-scored N3 sleep predicted neurobehavioral vigor, establishing a benchmark against the current standard. Regression models revealed no significant association between N3 and PVT-based vigor across all conditions tested. Specifically, four sets of regressions — comparing N3 fraction during the baseline sleep satiation night (SLEEPSAT3) with performance during baseline, resilience, and recovery wakefulness periods, and N3 during the first recovery sleep night (SLEEPR1) with next-day recovery performance — all yielded non-significant results (p = 0.4–1.0). Repeating these analyses using total N3 duration yielded equivalent null results, as did a normalized measure of recovery night N3 relative to satiation night N3. To confirm the null results were not attributable to failed experimental manipulation, we verified that N3, sleep onset latency, sleep efficiency, and PVT-based vigor all changed as expected across sleep pressure conditions (Figures S1, S2). Across multiple operationalizations of the deep sleep variable and multiple sleep pressure conditions, clinically-scored N3 carried no detectable functional signal for neurobehavioral resilience to, or recovery from, extreme sleep loss.

### Sleep Satiation SWA-Based Deep Sleep Negatively Predicts Vigor — Evidence for Residual Sleep Debt

Based on the hypothesis that clinical staging discards functionally relevant signal, we examined whether SWA power (1–4 Hz) during sleep provided greater predictive resolution than N3 scoring. Sleep-satiation baseline SWA (SLEEPSAT3) showed a consistent negative association with subsequent vigor — trending for resilience (p = 0.08) and reaching significance for recovery (p = 0.03) (Figure 1). We additionally examined an adjusted SLEEPSAT3 SWA — normalized by dividing by SLEEPR1 (recovery sleep night) SWA to account for inter-individual differences in baseline SWA amplitude — which showed a consistent negative correlation with recovery at a comparable effect size, providing convergent evidence for the unadjusted result. SLEEPR1 alone failed to show a significant relationship. We revisit this later. Together these findings are consistent with the interpretation that sleep-saturated SWA reflects residual sleep debt rather than enhanced recovery — higher SWA at baseline indexing greater homeostatic pressure, and thus predicting poorer subsequent vigor.

**Figure 1.**
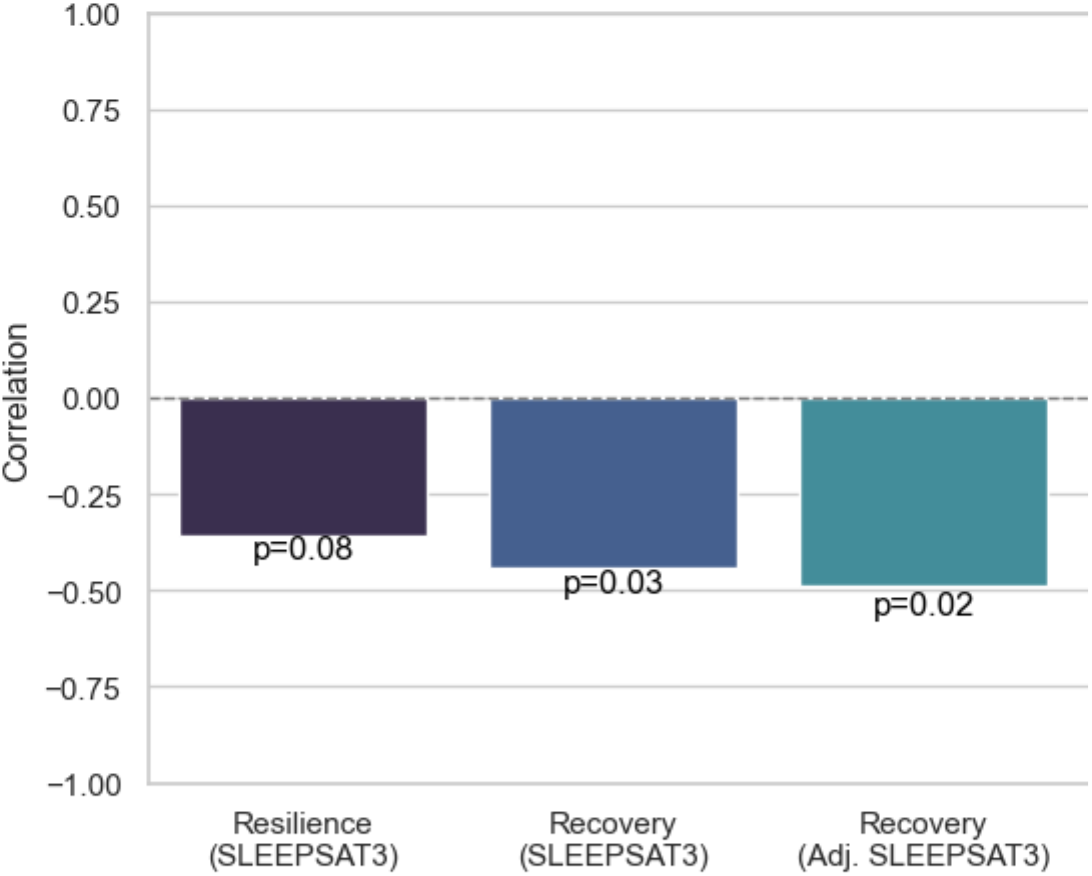
Inter-individual differences in resilience and recovery vs SWA during the third satiation sleep period. N=23.

### Wakefulness SWA During Sleep Deprivation Positively Predicts Vigor — Evidence for Active Recovery

Having established that sleep-satiation SWA negatively predicts vigor, consistent with residual debt, and that recovery sleep SWA carried little independent signal, we next asked whether SWA measured during the sleep deprivation wakefulness period (WAKESD) carried additional information. Unadjusted WAKESD SWA showed a significant positive association with recovery (r = 0.40, p = 0.046) but fell short of significance for resilience (r = 0.30, p = 0.1). The consistent positive direction across both outcomes was suggestive of a genuine recovery effect. Recognizing that the normalization denominator may confound interpretation, we examined whether normalizing for inter-individual differences in SWA amplitude clarified the signal. An adjusted WAKESD SWA — normalized by dividing by SLEEPSAT3 SWA — substantially strengthened both associations: explaining 38% of variance in resilience (r = 0.62, p = 0.001) and 61% in recovery (r = 0.78, p = 7×10^−6^) (Figures 2a, 2b). Critically, this association was positive — the opposite sign from the sleep satiation result — consistent with the interpretation that wakefulness SWA under sleep pressure reflects active neural recovery rather than residual debt. In the following section, we provide strong evidence that this is an independent sleep-like recovery effect.

**Figure 2.**
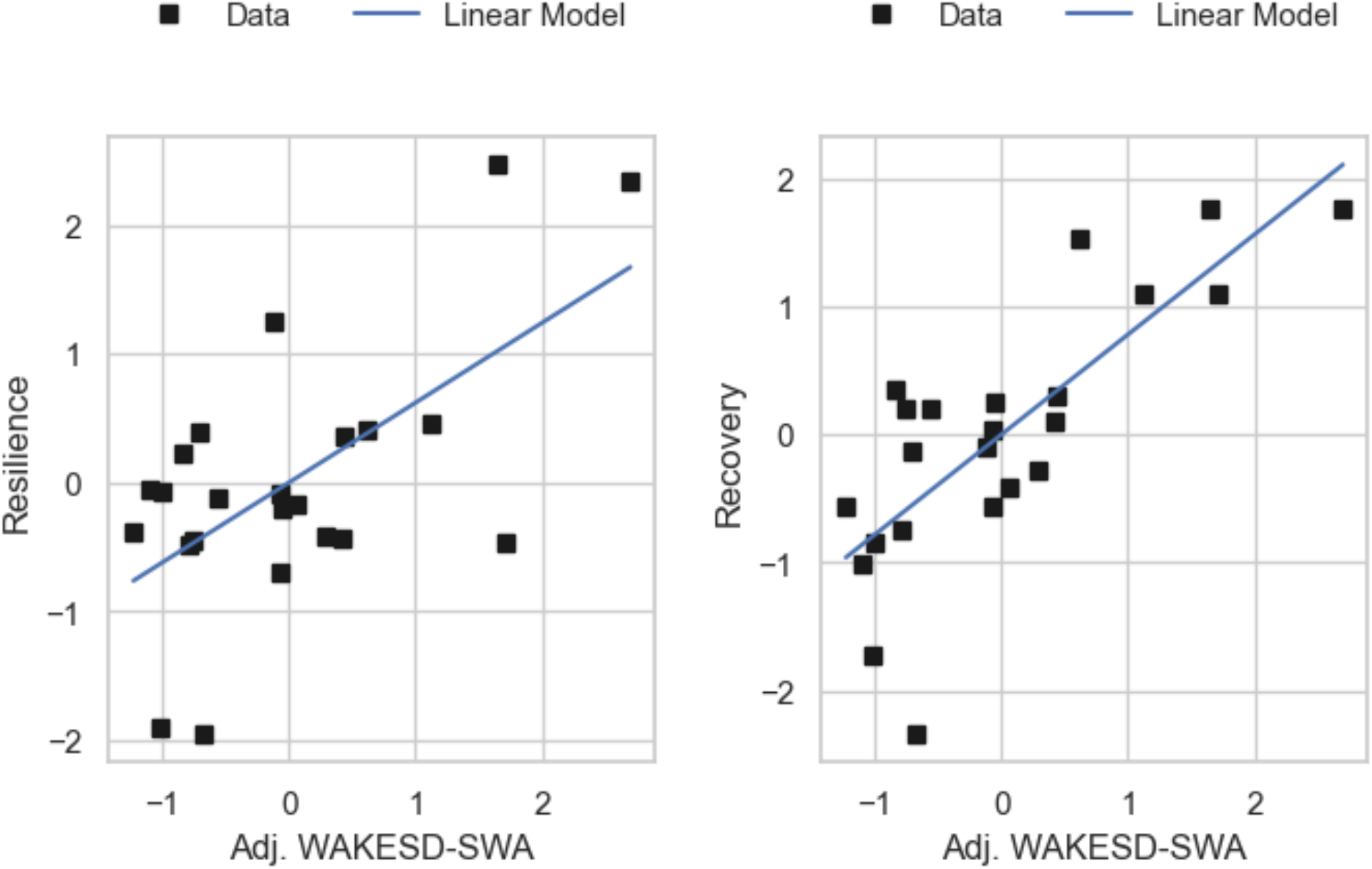
Inter-individual differences in resilience and recovery vs adjusted SWA during the sleep deprivation period. **A)** Resilience vs sleep deprivation period wake SWA (r=0.62, p=0.001, N=23). **B)** Recovery vs sleep deprivation period wake SWA (r=0.78, p=7×10^−6^, N=23).

Furthermore, EEG data were collected during Maintenance of Wakefulness Trials (MWTs) and performance testing outside of these, with individuals under strict observation controlling for frank sleep. We confirmed that these effects survived re-analysis with sleep scoring and removal of any scored-sleep epochs (see Supplementary Materials). The staggered EEG and performance testing also motivated the hypothesis that individuals with higher wakefulness SWA are more resilient in performance, for which we provide preliminary supporting evidence in the Supplementary Materials.

### Deep Sleep Effects Are Context-Dependent: Resolving the SWA Paradox

Thus far, univariate models have revealed context-dependent associations — sleep satiation SWA negatively predicting vigor and wakefulness SWA positively predicting vigor — that together constitute the Deep Sleep Dual Measurement and Causal Indeterminacy Problem. To test whether these opposing signals reflect independent sources of variance, we entered SWA from multiple time periods simultaneously into multivariate regression models (Figure 3). Both effects survived: sleep satiation SWA retained its negative association, indexing residual debt, while wakefulness SWA retained its positive association, indexing active recovery. The strongest and simplest model regressed recovery on SLEEPSAT3 and WAKESD SWA simultaneously (R^2^ = 0.54, standardized betas of -0.5 and +0.5 respectively, p = 0.001 for both). Furthermore, the interpretation of recovery night SWA depended entirely on model specification — when sleep satiation SWA was included, recovery night SWA carried a positive association; when excluded, it absorbed the debt signal and turned negative. The same measure, same recording period, opposite interpretation depending on temporal context. This is direct empirical confirmation of the Deep Sleep Dual Measurement and Causal Indeterminacy Problem.

**Figure 3.**
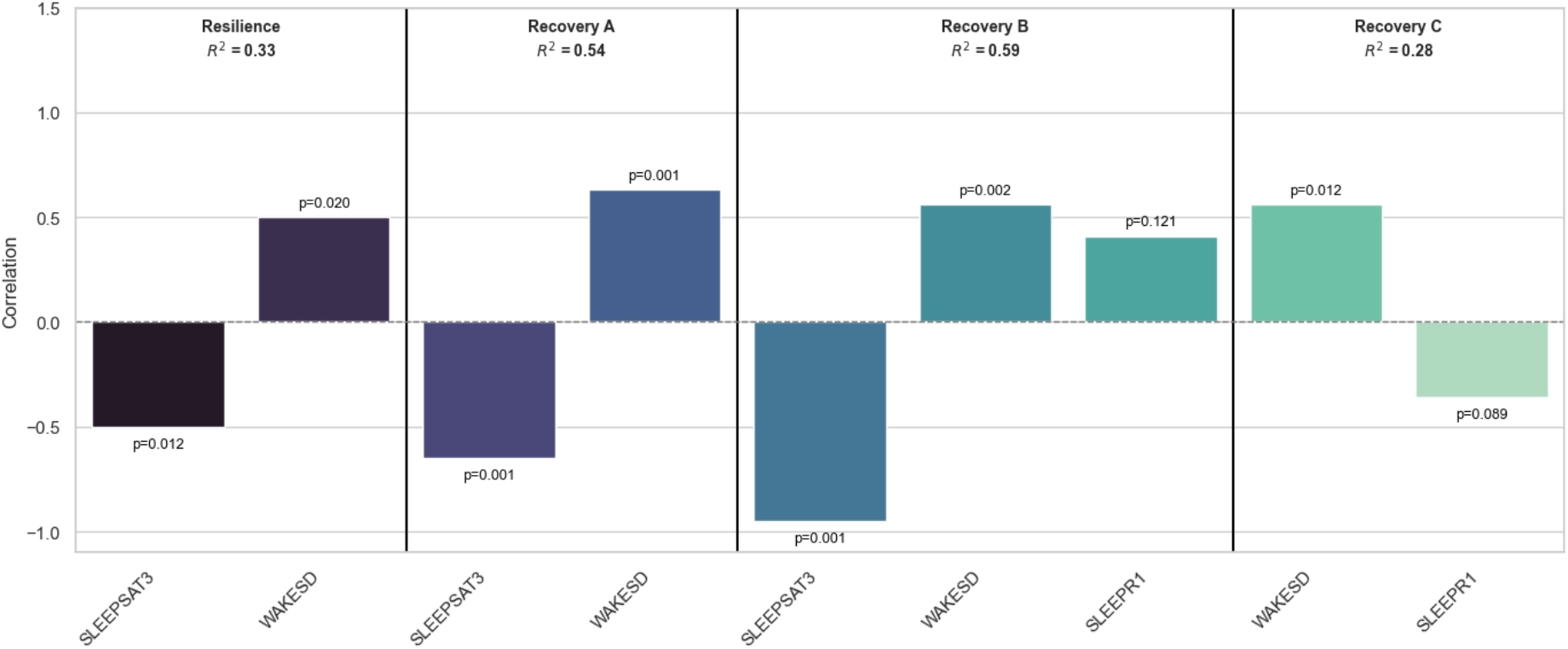
Multivariate resilience and recovery models. N=23.

## Discussion

In this study, we demonstrate that deep sleep measures reflect both ongoing recovery from neurobehavioral fatigue and accumulated sleep debt — a duality that is precisely what the Deep Sleep Dual Measurement and Causal Indeterminacy Problem predicts. Without proper contextualization, endogenous changes in these physiological markers cannot reliably be used to infer functional states or outcomes. While SWA measures showed strong associations with fatigue, clinical scoring of N3 provided no predictive value — a finding anticipated by Borbély himself, who characterized NREM staging as ‘a crude and rather arbitrary subdivision of a continuous process’^1^.

The discrepancy between SWA and clinically scored deep sleep may seem at odds with large population studies like All of Us, which report significant correlations between consumer-derived sleep staging and chronic illness traits^4^. However, these associations warrant careful interpretation. Large sample sizes alone cannot disambiguate causal indeterminacy — they simply render detectable whatever directional bias happens to dominate in that particular dataset. This problem is compounded by consumer devices, which do not directly measure neurophysiology — their algorithms infer sleep architecture from peripheral correlates that are nonspecific to the underlying electrophysiology and unpredictably confounded by the very conditions being predicted, and their fidelity degrades when applied to the clinical populations whose outcomes are being predicted^26-28^. These interpretation issues extend to direct SWA measurement as well — della Monica et al.^11^ found that SWA% reversed sign as a predictor of subjective sleep quality across age groups, the same problem of sample composition determining the direction of association.

Our results also shed light on resilience to sleep loss. Many practices — mindfulness, meditation, rest breaks, stress reduction — are explored as ways to mitigate cognitive load during wakefulness, yet their neurophysiological mechanisms remain poorly understood. Our findings suggest that covert sleep-like neural activity during wakefulness, indexed by wake SWA, may represent one such mechanism — and that individual differences in resilience may reflect the degree to which the brain engages sleep-like processes while behaviorally awake. This reframes a foundational assumption of the two-process model: Process S may not accumulate monotonically as a function of time awake but may be partially discharged in real time through covert neural activity. Put simply, the brain does not wait passively for behavioral sleep to repay its debt; it steals rest where it can. This also resolves a puzzle in prior resilience modeling work. Counterintuitively, Subramaniyan et al. (2024)^25^ found that higher baseline sleep SWA — ostensibly a marker of deeper, more restorative sleep — predicted greater vulnerability to sleep deprivation rather than greater resilience. Yet extracting this signal required substantial methodological scaffolding and still yielded only moderate performance (AUC = 0.68, sensitivity = 0.50). Both the counterintuitive direction and the methodological difficulty are explained by the same thing: baseline sleep SWA indexes homeostatic debt rather than recovery capacity, and trying to build a classifier on one half of a constitutively context-dependent signal is precisely as hard as the results suggest. The Dual Measurement and Causal Indeterminacy Problem provides the first unified account of these contradictions.

These findings call for a fundamental reappraisal of how deep sleep is measured, modeled, and interpreted. Clinical staging should be treated as descriptive architecture rather than a functional biomarker. Consumer sleep scores and the AI coaching pipelines built on them lack the temporal modeling and physiological grounding necessary for valid functional inference — and the compounding layers of measurement, causal, and device-level indeterminacy make them particularly unreliable. Statistical power alone is not a substitute for construct validity, and these limitations should be confronted before clinical adoption and, more urgently, before millions of users act on them. The Dual Indeterminacy framework provides a principled basis for next-generation clinical sleep metrics — physiologically grounded, temporally contextualized, and validated against functional outcomes rather than against staging categories that themselves lack predictive validity^29^.

## Methods

### Data

Participants were 23 healthy young adults screened for sleep disorders and health problems. To help ensure that participants entered the laboratory with minimal sleep debt, participants were asked to maintain an 8-hour sleep schedule while at-home for at least 7 days. This was followed by three baseline nights in the sleep laboratory with 10-hour time-in-bed (TIB) each night. Fatigue was assessed using objective and subjective scores. The former was assessed using the neurobehavioral gold-standard, reaction time on visual computer-based 10-minute psychomotor vigilance task (PVT)^21^. Testing was conducted at regular 75-minute intervals (interstimulus interval [ISI]: 2-10 seconds, with a uniform distribution).

### Study Protocol

In brief, the sleep protocol consisted of establishing a sleep satiated (rested) state; then, after a full normal wake period, a short sleep opportunity with or without SO stimulation; followed by sleep deprivation; and, finally, recovery sleep periods. We used five time periods defined by the study protocol (see Figure M1) for predictor and outcome measures: 1) the sleep satiation night prior to sleep deprivation (SLEEPSAT3), 2) the wake period immediately following sleep satiation (WAKESAT3)(aka baseline), 3) the sleep deprivation wake period (SD), 4) the first sleep recovery night (SLEEPR1), and 5) the wake period after the first recovery night (WAKER1). Our SWA power measures were intentionally agnostic to sleep-wake behavior and sleep-scoring. We visually confirmed that these putative SWA predictive signals were likely due to DOWN-state events (the critical component of SWA)^22’23^.

**Figure M1.**
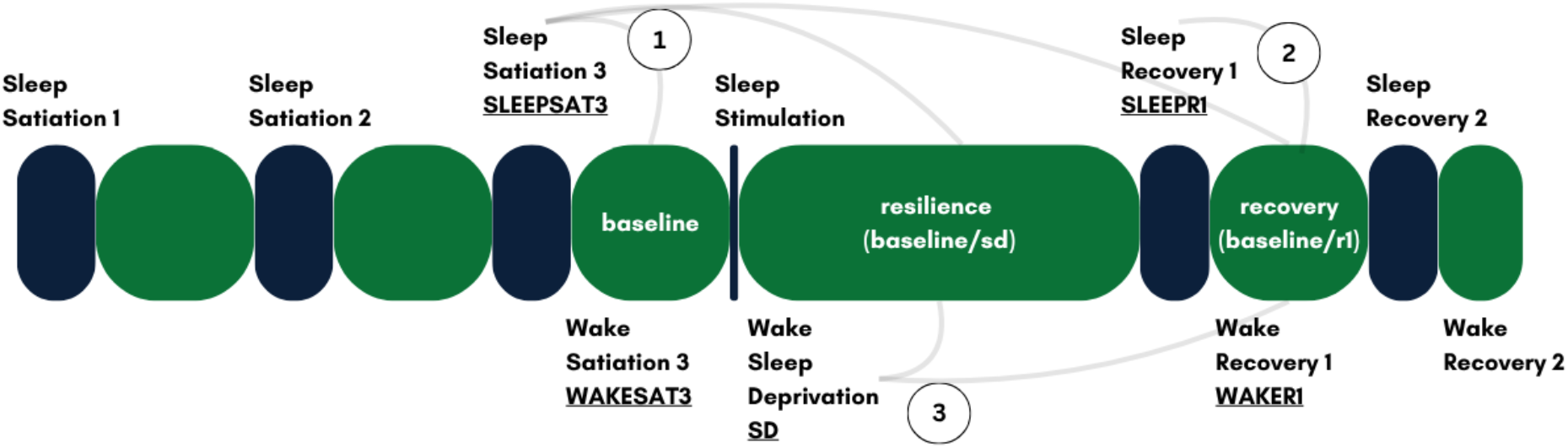
Sleep study protocol, measures, and regressions. Illustration of the experimental protocol (blue=scheduled sleep, green=scheduled wake). Starts with 8-hour scheduled sleep at home (not shown), followed by laboratory study days beginning with three sleep satiation nights. This was followed by a 17-hour wake period (WAKESAT3), a brief sleep opportunity, sleep deprivation (45 hours), and two nighttime recovery periods.

### Polysomnography Preprocessing

To obtain high-quality artifact free samples, including removal of eye-blinks and movement that can confound slow wave activity detection (during wake), we performed several preprocessing steps focused on 30-second EEG epochs. This workflow is illustrated below:

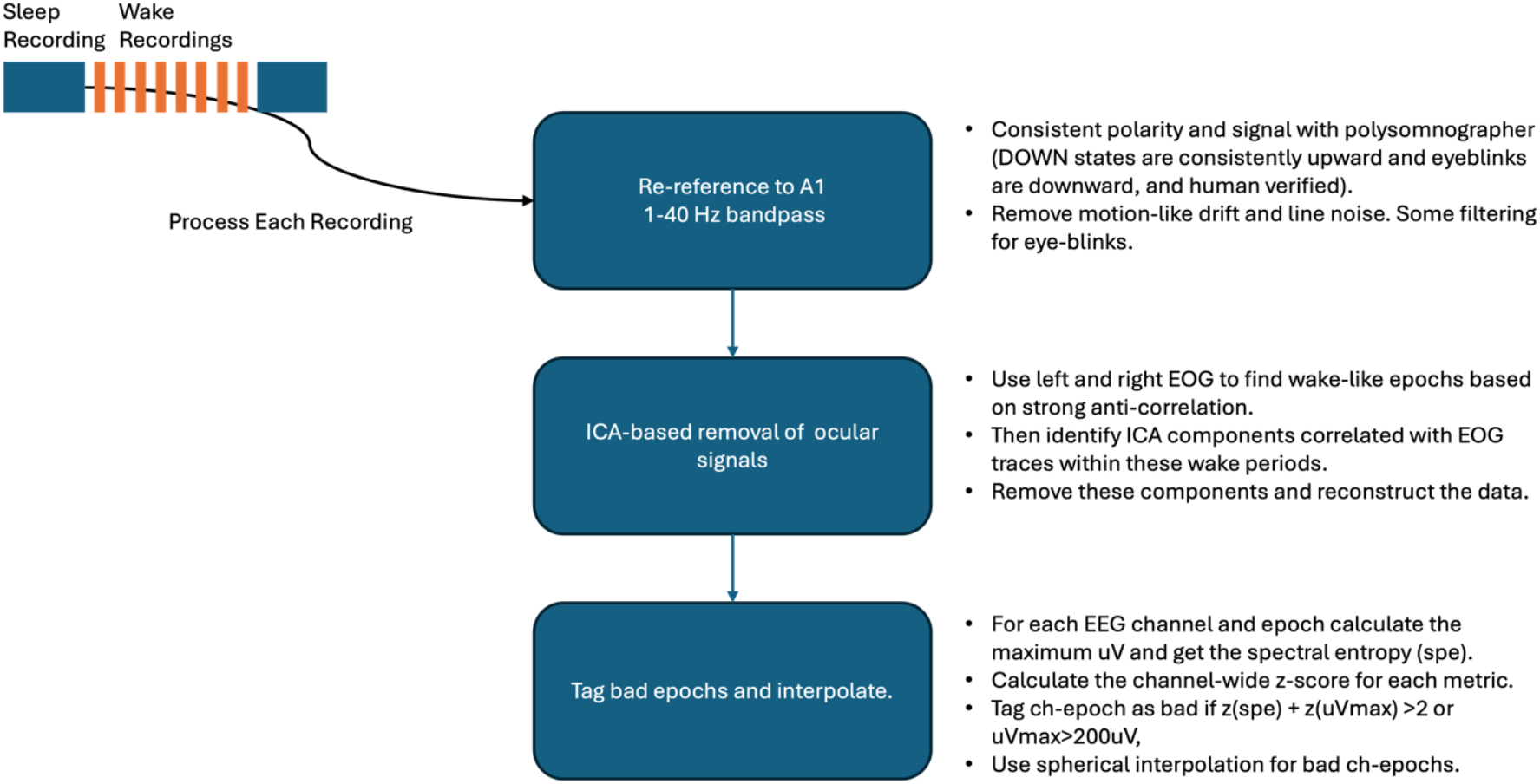

### PVT Processing

Our main outcomes were defined by two measures of vigor (inverse of fatigue) across individuals - resilience to and recovery from extreme sleep loss. Vigor (fatigue) was defined using objective PVT performance. For both sleep predictors and vigor (or fatigue) outcomes, we used time-weighted average estimates for given time periods, a form of interpolation. This was derived by taking the discrete integral across the period and computing the weighted average, with weights determined by the duration between samples. This is a commonly-used method for interpolating irregularly sampled data. It is intended to give equal weight to all samples. For example, this was important for the wakefulness metrics derived from the MWTs because the samples become more widely distributed as sleep debt accrues across individuals; less wake data with sleep deprivation. We also had irregular sampling due to quality control measures that eliminated some epochs, including during continuous recordings. For consistency, time-weighted averaging was applied to all measures.

We defined resilience as the ratio between time-weighted average PVT-reaction time (PVT-RT) during WAKESAT3 over its counterpart during SD. Thus, a larger value means a more rested state for that individual – with a value of one being equivalent to their post-sleep satiation (baseline) state and a value higher than one being superior. (In one analysis, we provide the inverse of this measure to reflect vulnerability, with one still indicating equivalence but greater than one indicating increased fatigue.) Recovery from extreme sleep loss was calculated as the ratio of the time-weighted average WAKESAT3 PVT-RT over WAKER1 PVT-RT, so higher values indicated greater recovery.

## Glossary

Neurobehavioral Fatigue: Objective fatigue measured using the Psychomotor Vigilance Test (PVT).
PVT (Psychomotor Vigilance Test): A neurobehavioral measure used to assess objective fatigue.
Recovery (from extreme sleep loss): A measure of vigor, calculated as the ratio of the time-weighted average PVT-RT during WAKESAT3 over the PVT-RT during WAKER1. Higher values indicate greater recovery.
Resilience (to sleep loss): A measure of vigor, calculated as the ratio of the time-weighted average PVT-RT during WAKESAT3 over its counterpart during SD. A larger value indicates a more rested state.
SWA (Slow-Wave Activity): Power in the slow-wave activity band (1-4Hz), regarded as the primary restorative component of deep sleep.
Vigor: The inverse of fatigue, defined using objective performance on the PVT through the measures of Resilience and Recovery.
SLEEPSAT3 (Sleep Satiation Night 3): The third night of 10-hour scheduled sleep, used for baseline physiological measures.
SLEEPR1 (Sleep Recovery Night 1): The first 8-hour recovery sleep period following the 45-hour sleep deprivation period.
WAKESAT3 (Sleep Satiation Wake Period): The 17-hour wake period immediately following sleep satiation, used as the baseline performance period.
WAKESD (Sleep Deprivation Wake Period): The 45-hour period of continuous wakefulness.
WAKER1 (Recovery Wake Period): The wake period after the first recovery night.

## Supplementary Materials

### S1. Fatigue (Vigor) Changes with Sleep Loss and After Recovery Sleep

As expected, vigor (the inverse of fatigue) was lower during sleep deprivation (the resilience test period) than after recovery sleep (Figure S1). This relative difference was more marked when normalizing by individual baseline PVT performance. In the unnormalized group analysis (Figure S1a), there was a trend difference between conditions (t = 1.6, p = 0.1). This became significant with normalization (Figure S1b), with slope β = 1.1 (p < 0.0001, R^2^ = 0.6). The coefficient greater than one is consistent with the grouped results in Figure S1a, confirming that vigor improved following recovery sleep.

**Figure S1.**
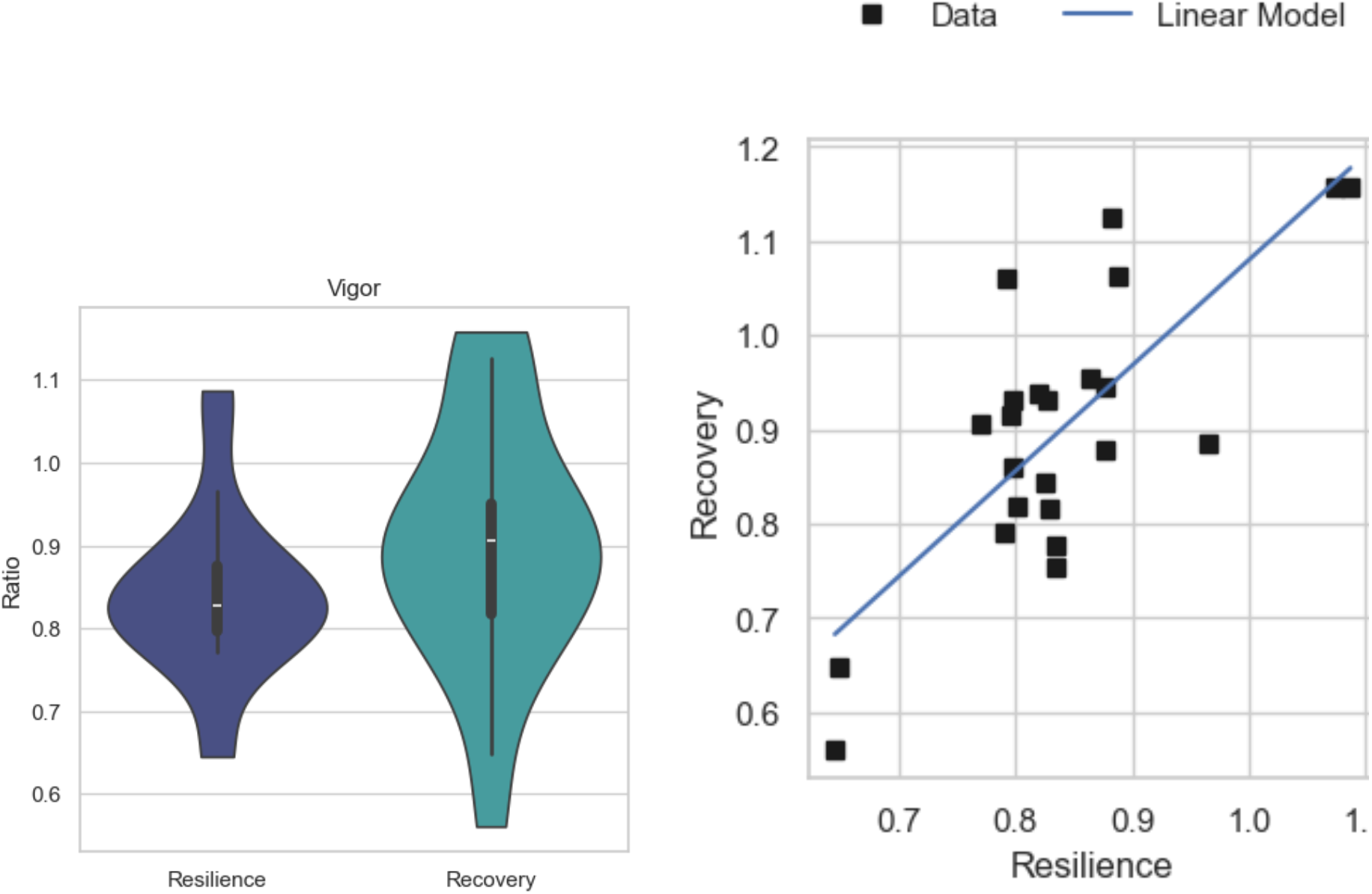
Fatigue (Vigor) Measures Across Wake Periods. (a) Group-wise variation in vigor during the sleep deprivation resilience phase and the recovery phase (t = 1.6, p = 0.1). (b) Regression of recovery vigor on resilience vigor (β = 1.1, p < 0.0001, R^2^ = 0.6). N = 23.

### S2. Marked Differences in Clinically-Scored Deep Sleep During Sleep Satiation vs. Recovery Sleep

We assessed the statistical distribution of sleep parameters for the baseline sleep satiation night (SLEEPSAT3) vs. the first recovery sleep night (SLEEPR1) for descriptive purposes; no formal statistical testing was conducted. As expected, recovery sleep was characterized by shorter sleep onset latency (SOL) and higher sleep efficiency (SE) (Figure S2). These findings also highlight that sleep history and context are critical for interpreting standard metrics — the sleep-satiated SOL and SE distributions overlapped with insomnia ranges, while recovery SOL fell almost entirely within the excessive sleepiness range. Both reflect normal, expected patterns for these conditions but can be misinterpreted without contextual framing. Furthermore, clinically-scored N3 increased during recovery, consistent with prior findings and indicative of a strong homeostatic response to accumulated sleep debt. The observed changes in sleep architecture and fatigue across sleep debt conditions were thus clear and expected. The following sections examine whether these clinical metrics predict functional states.

**Figure S2.**
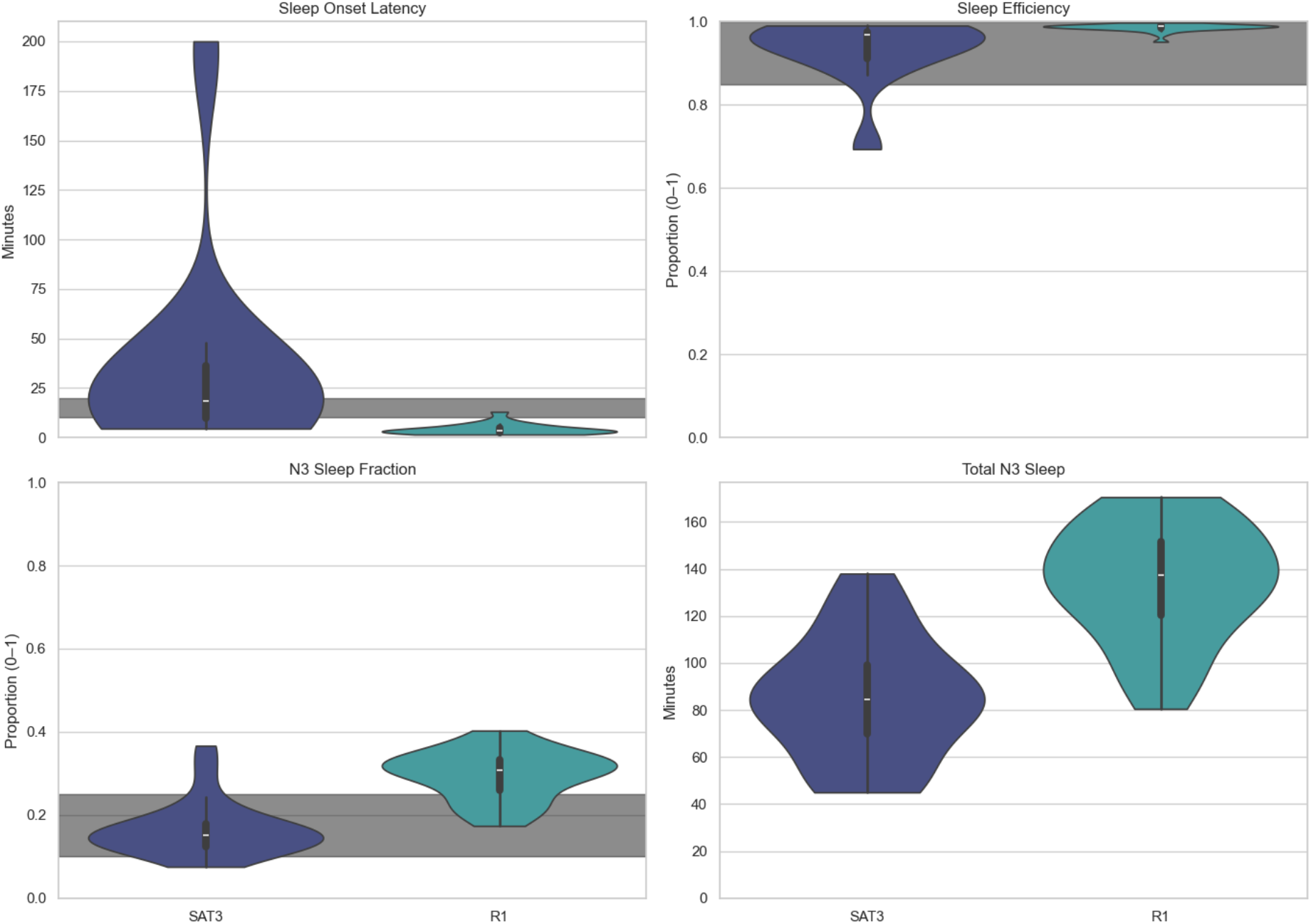
Sleep onset latency, sleep efficiency, and clinically-scored N3 during sleep satiation (SLEEPSAT3) vs. recovery sleep (SLEEPR1). Dark gray band indicates the typical healthy sleep range. Shown for illustrative purposes. N = 23.

### S3. Sleep Deprivation Wakefulness SWA Effect Holds Under Conservative Scoring

A potential concern is that subjects may have been experiencing frank sleep during the sleep deprivation recording period. However, recordings were taken during monitored Maintenance of Wakefulness Tests (MWTs) with strict instructions to terminate recordings upon any detected sleep. To further test robustness, we selected the subset of subjects with scored MWT data (n = 16) and removed any scored sleep epochs prior to analysis. The resilience and recovery correlations remained significant: r = 0.7 (p = 0.003) and r = 0.8 (p = 3×10^−5^), respectively, confirming that the wakefulness SWA effect is not attributable to covert sleep during the recording period.

### S4. Rate of Fatigue Accumulation Varies Across Individuals and Is Predicted by Wakefulness SWA

Previously reported results for sleep deprivation physiology consistently show a positive association between SWA and fatigue at the group level — the opposite direction from our cross-individual finding, where elevated wakefulness SWA was associated with greater vigor. This apparent conflict is resolved by distinguishing group-averaged from cross-individual effects.

To illustrate this, we first examined the association between SWA and vulnerability (the inverse of our resilience measure) at the group level. As shown in Figure S3a, elevated SWA was associated with increased vulnerability to sleep loss (r = 0.76, p < 0.001), consistent with prior literature — on average across individuals, SWA and fatigue increase together as sleep debt accumulates, with a circadian modulation also evident in both traces.

This group-vs-individual dissociation motivates a specific hypothesis: while all individuals are affected by prolonged wakefulness, individuals differ substantially in their relative vulnerability, and this variability may reflect individual differences in a protective covert wakefulness SWA. Examination of this hypothesis confirmed the prediction (Figure S3b) — individuals in the lowest vulnerability tertile showed higher wakefulness SWA than those in the highest vulnerability tertile, and this difference became more pronounced with greater sleep loss. As noted, wake EEG recordings were collected separately from performance testing; whether elevated SWA also occurred during task execution remains unknown. The results suggest this may not be the case, and continuous recordings during performance testing are warranted to investigate this further.

**Figure S3.**
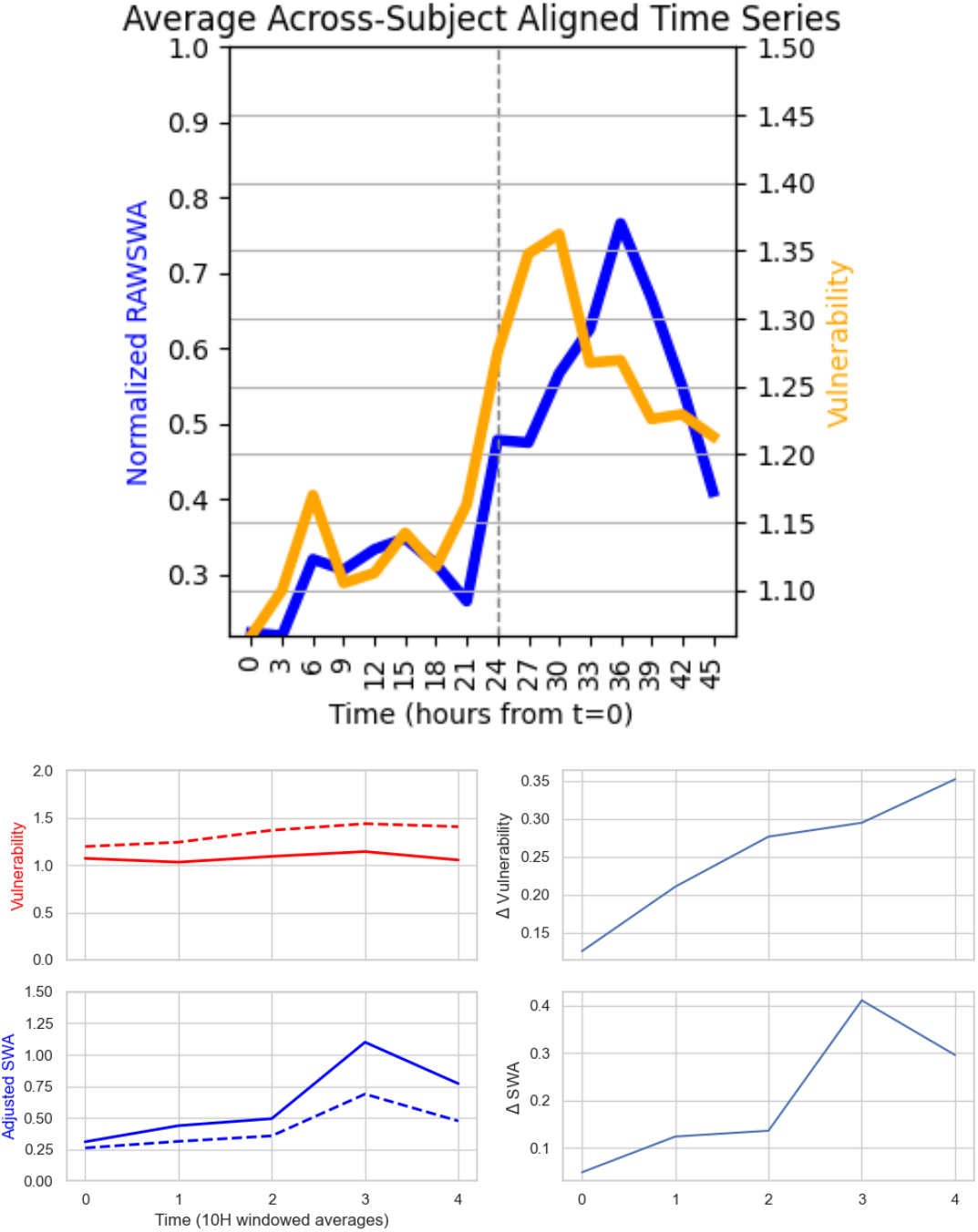
Vulnerability to sleep loss and adjusted SWA during the sleep deprivation period. (a) Group-average SWA and vulnerability traces across the sleep deprivation period. (b) Left panel: average traces for the top (dashed) and bottom (solid) vulnerability tertiles. Right panel: difference in vulnerability and SWA between the top and bottom vulnerability tertiles. N = 23.

